# Multigenerational effects of tire wear particle leachate through trophic transfer from producer to primary consumer

**DOI:** 10.1101/2023.07.08.548196

**Authors:** Yanchao Chai, Haiqing Wang, Jiaxin Yang

**Author notes:** Equal contributions to this article (Yanchao Chai, Haiqing Wang).

## Abstract

Tire wear particle (TWP) and their leachate have been known toxic for aquatic organism due to additives released from the rubber matrix. However, it is not clear whether the ecotoxicity of TWP leachate could be transferred through algae-zooplankton food chain especially after multi generations. In this work, the effect of different concentrations TWP leachate on growth of microalgae, *Chlorella pyrenoidesa*, was studied. Subsequently, those algae were fed to zooplankton rotifer, *Brachionus calyciflorus*, which continuously lasted for ten generations to explore multigenerational accumulation of TWP leachate ecotoxicity through food-chain transfer. The results showed that the TWP leachate displayed growth inhibition for algae with evident concentration effect. For rotifer fed with those contaminated microalgae, though the first two generations showed hormesis, the ecotoxicity exhibited after 2 generations, characterized by reduction of lifespan and offspring number. The ecological effects of TWP leachate were transferred from algae to rotifer. In addition, the ecotoxicity gradually aggravated along with generation passage and exposure concentration of algae. What was even more, population passage of rotifer collapsed totally with no offspring after 5-generations feed by algae exposed to high concentration TWP leachate. Based on those, it is summarized that the ecological effects of TWP leachate indeed can be transferred from low to high trophic level in food chain, and accumulate across generations passage. The indirect non-contact exposure through food chain should be considered at the risk assessment of TWP. Single generation exposure will underestimate their ecological risk from long term.

## 1. Introduction

Sporadic collective deaths of salmon in urban stream have been linked to the extractable additive from tire wear particle (TWP) in road runoff (Tian et al., 2021), which should be a wake-up call about the effects of TWP on aquatic ecosystem function and stability. As increasing of tire utilization and non-point source pollution without effective control, more TWP is accessible to aquatic ecosystem through runoff and atmospheric sedimentation. Once into water, TWP would release complex toxic additives as leachate, which imposed impacts on aquatic biome (Halsband et al., 2020; Knight et al., 2020; Kole et al., 2017; Wagner et al., 2018). However, current studies of these impacts have primarily focused on the effects of TWP leachate at just single specie and generation. It has not been reported whether the ecological effects of TWP leachate could be transferred through food-chain among different species, and how this trophic transfer respond along with multigenerational exposure.

It has been well known that there are trophic transfer and multigenerational effects for many typical environmental pollutions, as they can accumulate *in vivo* and be passed to higher consumer, known as bioaccumulation, and aggregate damages after continuous exposure of multiple generations (Guo et al., 2012; Stuligross and Williams, 2021). TWP leachate has been shown toxic effects on different trophic-level taxa (algae, zooplankton, fish) in direct exposure to single specie(Marwood et al., 2011), but food-chain transfer of ecotoxicity need to be verified in further. According to author’s previous study, direct exposure of TWP leachate across 7 generations to rotifer exhibited multigenerational accumulation effects (preprint, DOI: 10.1101/2022.10.27.513999). It needs further study to explore whether the indirect exposure by trophic transfer show similar multigenerational effects. Those delayed impacts can be ignored in current exposure on single generation, which underestimates effects of stressors.

We investigated the effect of different concentrations TWP leachate on growth of microalgae, *Chlorella pyrenoidesa*. Subsequently, those algae were fed to zooplankton rotifer, *Brachionus calyciflorus*, which continuously lasted for ten generations to explore multigenerational accumulation of TWP leachate ecotoxicity through food-chain transfer. This study will provide more precise risk assessment for TWP based on indirect non-contact trophic relationship and long term exposure.

## 2. Materials and Methods

### 2.1 Preparation of TWP leachate and rotifer

TWP were purchased from a factory that recycles and smashes scrap tires into particles as artificial turf material. These TWP were sieved through 100 μm nylon net, then sealed and stored at 4°C. TWP were soaked with hard synthetic freshwater (96 mg NaHCO_3_, 60 mg CaSO_4_, 60 mg MgSO_4_ and 4 mg KCl in 1 L distilled water) (USEPA, 1985) at 2500 mg/L in glass bottle and mixed uniformly. The mixture was aerated continuously for 15 days in dark condition at 25°C. It has been demonstrated that there is equilibrium between water and TWP for most components after 14-days leaching (Capolupo et al., 2020; Selbes et al., 2015). TWP leachate mother liquor was obtained by sieving the mixture through 38 μm nylon net, and frozen at -20°C for temporary storage.

Rotifer *B. calyciflorus* clone strain was established from resting egg that was presented by Professor T.W. Snell of Georgia Institute of Technology, USA. The above hard synthetic freshwater was used as culture medium. The culture condition was kept in illumination incubator with 16L:8D light schedule with light intensity of 4000 lux at 25°C. Rotifers were fed with 5×10^6^ cell/ml *C. pyrenoidesa*, and the medium was replaced every day. Before experiment, some rotifer with amictic egg were chosen from clone strain and synchronized (Kaneko et al., 2016).

### 2.2 Concentration effect of TWP leachate on algae growth

Microalgae, *C. pyrenoidesa*, were separately inoculated into 1-L conical flask with 500ml BG-11 culture medium with different TWP leachate concentrations (0, 500, 1000, 1500, 2000, 2500 mg/L), which were prepared by diluting TWP leachate mother liquor. Every treatment included three replicates. The initial density of algae was 10^6^ cell/ml. Those algae were kept at uninterrupted illumination with light intensity of 4000 lux at 25°C. Air filtered through 0.22μm acetate fiber filter was pumped constantly into culture medium to prevent algae aggregation settlement and sustain carbon source level. The algae density was calculated by hemocytometer every day. The algae were harvested by centrifugation after 8 days cultivation, and stored at 4°C.

### 2.3 Trophic transfer of ecotoxicity from algae to rotifer across multi generations

After synchronization of parent, the rotifer individuals with amictic eggs were pick out. Then, neonate born within 4 hours were put into 24-well plate. There was one individual in one well with 1 ml hard synthetic freshwater, in which harvested algae exposed to TWP leachate (500, 1000 mg/L stands for low and high concentration, separately) were added at the density of 5×10^6^ cell/ml. The plates were placed in illumination incubator with 16L:8D light schedule with light intensity of 4000 lux at 25°C. Survival and offspring number were checked every 12 hours, neonates were taken out and transferred into corresponding new culture medium same with their parent. Each treatment in every generation included 48 individuals. The total 10 generations were fed and observed.

### 2.4 Statistics

The average individual reproduction and lifespan of rotifers at treatment were calculated by dividing the total offspring number and survival time by individual numbers (picking out mictic individuals). To eliminate intergenerational difference and display treatment effects more directly, within same generation, relative value to control group (%) of lifespan and reproduction in different treatments was utilized (Formula 1). Positive value means promotion effect, otherwise inhibition.

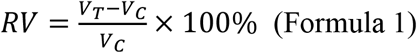

Where RV, V_T_ and V_C_ was relative value, treatment group value and control group value of average individual lifespan and offspring number, respectively.

## 3. Results and discussions

Compared to control group, the growth of *C. pyrenoidesa* exposed to TWP leachate were inhibited with evident concentration-dependent manner (Figure 1). This is not surprising as TWP leachate has been verified toxic to algae, which is attributed to toxic additives release from tire.

**Figure 1.**
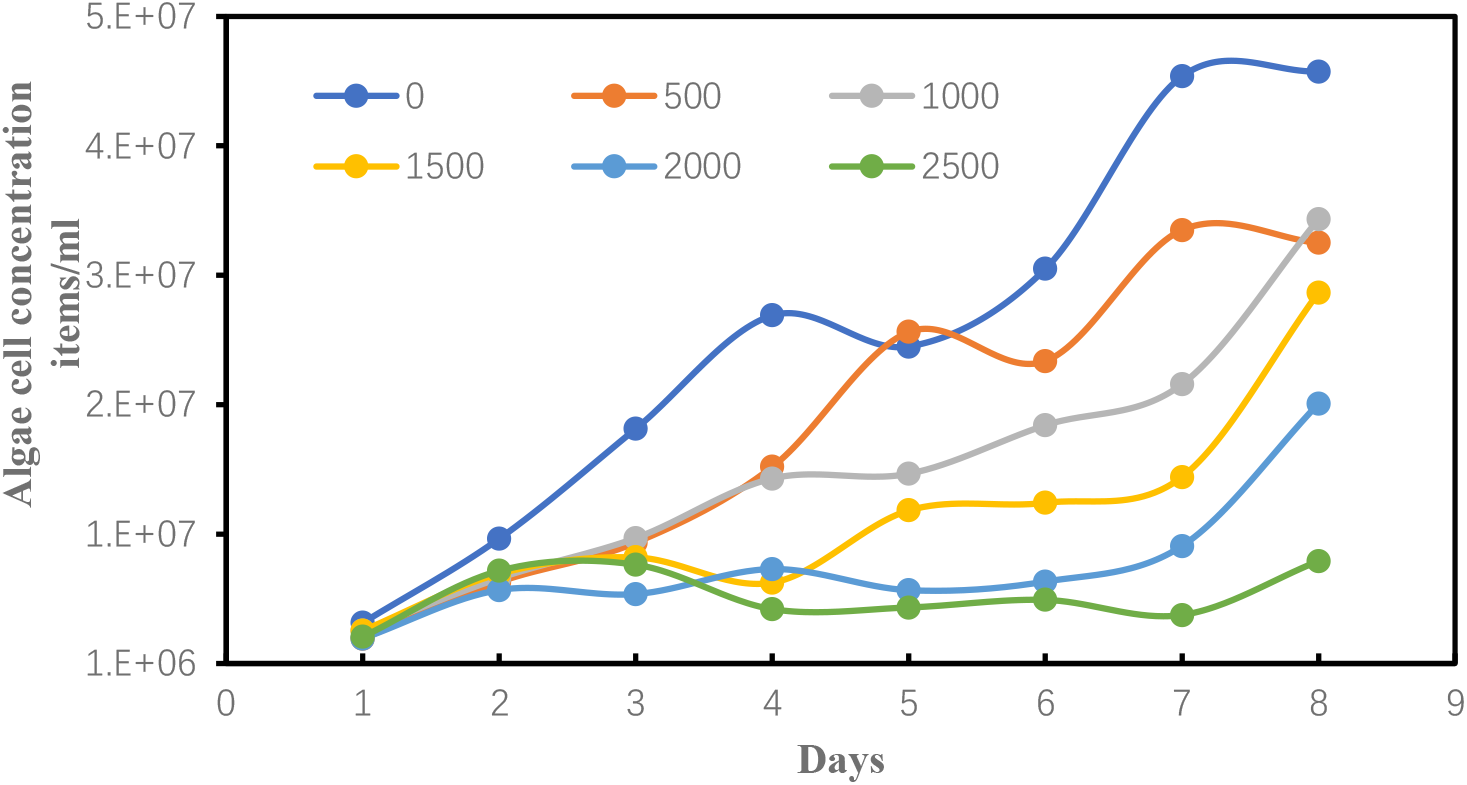
Growth curve of algae at different TWP leachate concentrations. Different color lines mean different concentration (mg/L).

When the rotifer were feed with algae exposed to TWP leachate (500 and 1000 mg/l were chosen to stand for low and high concentration), the average individual lifespan and offspring number were promoted in the first two generation (Figure 2). This phenomenon could be called as hormesis that is a central concept of toxicology to account for mild stress-induced beneficial effect and has been observe in many toxic substances (Sun et al., 2021). The hormesis was similarly observed in author’s previous study when rotifer were directly exposed to low-dosage TWP leachate (preprint, DOI: 10.1101/2022.10.27.513999). Analogy to direct contact, algae transferred this effect of TWP leachate to rotifer by non-contact trophic relationship.

**Figure 2.**
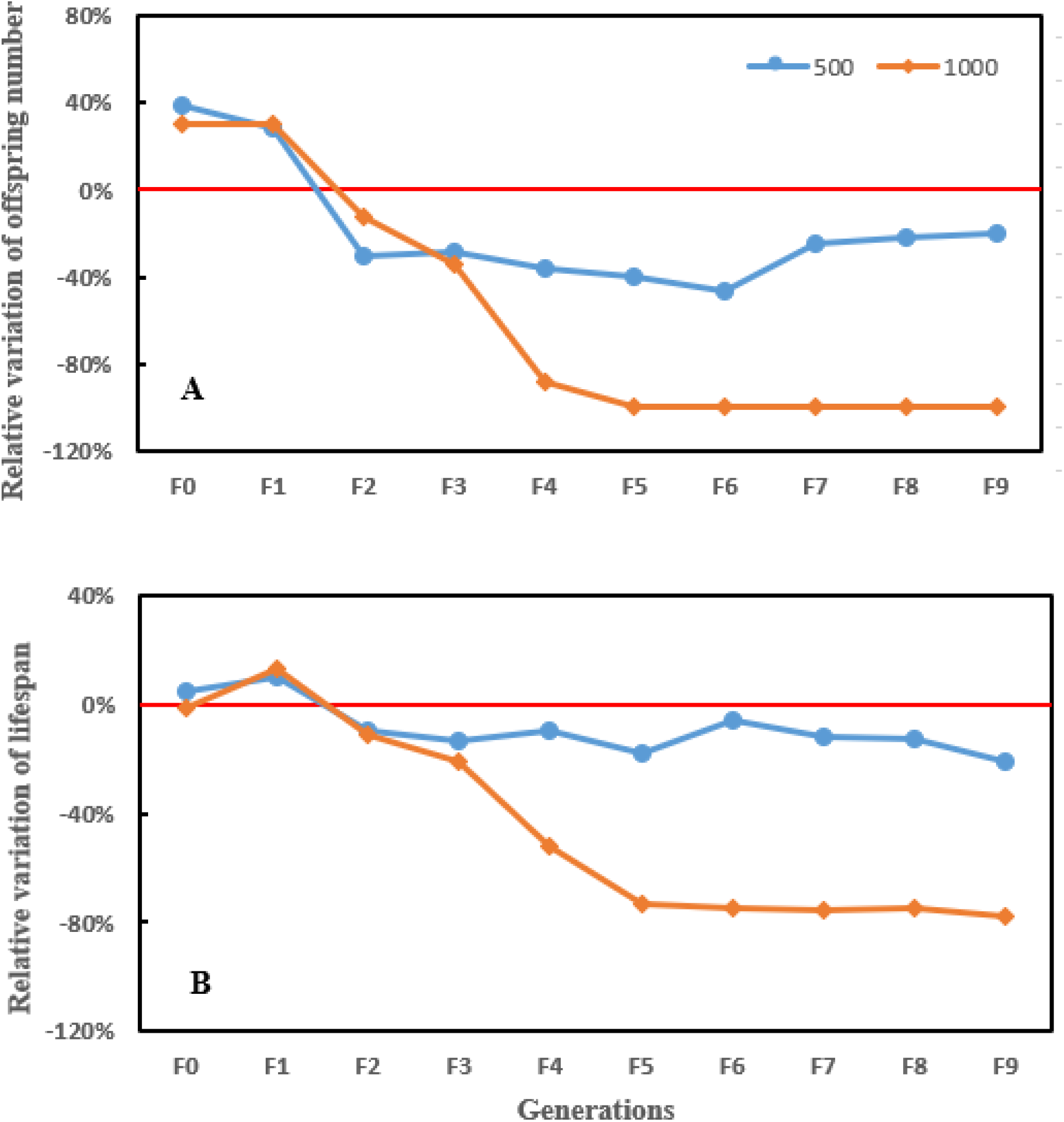
Relative value to control group for average individual lifespan (A) and reproduction (B) of rotifer fed with algae exposed to 500 and 1000 mg/l TWP leachate across 10 generations. Positive value means promotion effect, otherwise inhibition. Different color represents different concentration (mg/l), respectively.

After 2 generations, the average individual lifespan and offspring number were inhibited with concentration effect, and the inhibition exhibited aggravation trend along with generations passage (Figure 2). The algae exposed to high-dosage TWP even triggered population collapse. This is also similar with author’s previous study in which rotifer were directly exposed to TWP leachate across continuous 7 generations (preprint, DOI: 10.1101/2022.10.27.513999). The hormesis would be reversed once pollution dosage exceeds the critical value, which means that the ecotoxicity of TWP leachate transferred from low trophic level also can exhibit multigenerational accumulation.

## 4. Conclusions

The ecotoxicity of TWP leachate can be transferred from low to trophic level, which also accumulates across generation passage. The indirect non-contact exposure through food chain should be considered at risk assessment of TWP, similarly, single generation exposure will underestimate their ecological risk from long term. It is worth noting that some species in ecosystem, especially high trophic level, may face the risk of continuous population recession along with generation passage if environmental TWP accumulate without effective control.

## Acknowledgements

This work was supported by the Graduate Research and Innovation Project of Jiangsu Province (KYCX20_1190).

## Author contributions

**Yanchao Chai:** Conceptualization, Methodology, Writing - original draft, Funding acquisition. **Haiqing Wang:** Conceptualization, Writing - review & editing. **Jiaxin Yang:** Supervision, Writing - review & editing.

